# *Drosophila* egg-derived tyrosine phosphatase (EDTP): a novel target for improved survivorship to prolonged anoxia and cellular protein aggregates

**DOI:** 10.1101/279885

**Authors:** Chengfeng Xiao, Shuang Qiu, Xiao Li, Dan-Ju Luo, Gong-Ping Liu

**Author notes:** Corresponding authors (C.X.) and (G-P. L.).

## Abstract

*Drosophila* egg-derived tyrosine phosphatase (EDTP), a lipid phosphatase that removes 3-position phosphate at the inositol ring, has dual functions in the oogenesis and the muscle performance during adult stages. A mammalian homologous gene *MTMR14*, which encodes the myotubularin-related protein 14, negatively regulates autophagy. Mutation of *EDTP/MTMR14*, however, causes at least three deleterious consequences: (1) lethality in the early embryogenesis in *Drosophila*; (2) “jumpy” phenotype with apparently impaired motor functions; and (3) association with a rare genetic disorder called centronuclear myopathy. Here we show that flies carrying a heterozygous *EDTP* mutation had increased survivorship to prolonged anoxia; tissue-specific downregulation of *EDTP* in non-muscle tissues, particularly motoneurons, extended the lifespan; and tissue-specific downregulation of *EDTP* in motoneurons improved the survivorship to beta-amyloid peptides (Aβ42) and polyglutamine (polyQ) protein aggregates. MTMR14 expression was evident in the hippocampus and cortex in C57BL/6J and APP/PS1 mice. Compared with C57BL/6J mice, APP/PS1 mice had reduced MTMR14 in the cortex but not in the hippocampus. Hippocampal expression of MTMR14 was increased and plateaued at 9-17 months compared with 2-6 months in C57BL/6J mice. Aβ42 treatment increased the expression of MTMR14 in the primarily cultured hippocampal neurons of Sprague/Dawley rats and mouse Neuro2a neuroblasts. We demonstrated a novel approach of tissue-specific manipulation of the disease-associated gene *EDTP/MTMR14* for lifespan extension and the improvement of survivorship to cellular protein aggregates.

## Introduction

*Drosophila* egg-derived tyrosine phosphatase (EDTP) is a lipid phosphatase that removes 3-position phosphate at the inositol ring of phosphatidylinositol 3-phosphate (PtdIns3P) and phosphatidylinositol (3, 5)-bi-phosphate [PtdIns(3,5)P2] [1]. EDTP possesses a function opposite to Vps34, a sole class III phosphoinositide 3-kinase [2,3], in the regulation of the PtdIns3P pool. The most interesting characteristic of EDTP expression is that there are two peaks, one at oogenesis [4,5], and another at adult stages [6,7]. The transcription of *MTMR14*, a mouse homolog of *EDTP*, also shows a peak at day five of differentiation in C2C12 myoblasts, followed by a decline [1]. Human *MTMR14* transcripts are detectable at the ages between 19 and 69 years in the tested tissues, including heart, brain, placenta, lung, liver, skeletal muscle, kidney, and pancreas [1]. Levels of *MTMR14* are highly coincident with those of *Vps34* in the human brain, heart, and skeletal muscles [8], indicating a tightly regulated biological process which is associated with cellular PtdIns3P levels.

The decline of EDTP between early embryogenesis and young adult stages is accompanied with the *Drosophila* metamorphosis, a process requiring extensive autophagy and apoptosis for histolysis [9]. These observations suggest a role for EDTP in the regulation of autophagy. This is indeed supported by the findings that PtdIns3P stimulates autophagy in human HT-29 cells [10], that MTMR14 negatively controls the autophagosome formation and maturation in mammalian cells [11], and that the EDTP/MTMR14 inhibitor, AUTEN-99, activates autophagy in human HeLa cells and mouse tissues [12]. Interestingly, the latter group also shows that overexpression of a modified *EDTP* (by skipping the first exon) in the fat body antagonizes the effect of AUTEN-67 in inducing autophagy [13].

Despite the negative regulation of autophagy, the deletion of *Drosophila EDTP* is lethal during embryogenesis or in the first instar, and germline clones with a null *EDTP* allele fail to produce mature oocytes [5]. Homozygous flies carrying a hypomorphic *EDTP* allele are short-lived with impaired motor functions and reduced fecundity [14]. Additionally, muscles of the *MTMR14*-deficient mice have decreased force production, prolonged relaxation and exacerbated fatigue [15]. A human MTMR14 missense variant (R336Q) is associated with centronuclear myopathy, a rare genetic disorder with muscle weakness and wasting [1]. Therefore, the function of autophagy initiation is likely overwhelmed by the lethality or disease-causing effects due to a ubiquitous loss of *EDTP/MTMR14*.

There are advantages of autophagy in degrading and recycling disrupted organelles, long-lived proteins, and denatured protein aggregates [10,16]. A strategy to maximize the potential benefit of *EDTP* is to manipulate its downregulation in the favorable tissues while leaving the expression intact in oocytes and as well as in muscles at adult stages. This seems to be feasible by using the *Drosophila* Gal4/UAS expression system [17]. We thus hypothesized that selective downregulation of *EDTP* in non-muscle tissues, particularly the central nervous system, extends lifespan and improves the survivorship to cellular protein aggregates in *Drosophila*.

In the current study, we demonstrate that heterozygous *EDTP* mutants had improved survival to prolonged anoxia, a condition mimicking the extreme hypoxia that induces autophagy in mammalian and human cells [18]. We also show that selective downregulation of *EDTP* in the fly motoneurons extended lifespan and increased the survivorship to beta-amyloid peptides (Aβ42) or polyglutamine (polyQ) protein aggregates. The expression of MTMR14 in the hippocampus and cortex in mice was evident, promising a potential application by targeting *EDTP/MTMR14* for the treatment of neurodegeneration.

## Materials and methods

### Animals

Fly strains used in this study and their sources were: *EDTP*^*DJ694*^ (FBal0160586) and *w*^1118^ (L. Seroude laboratory, Queen’s University), D42-Gal4 (Bloomington Drosophila Stock Center (BDSC) #8816), UAS-EDTP-RNAi (II) (with an RNAi sequence CAGTAGTGTAATAGTAATCAA targeting exon 3 of *EDTP*, FBst0041633), UAS-EDTP-RNAi (III) (with an RNAi sequence CAGCTACGACGAAGTCATCAA targeting exon 2, FBst0036917), UAS-Aβ42 (BDSC #32038), UAS-Httex1-Q103-eGFP [19]. Flies were maintained with standard medium (cornmeal, agar, molasses, and yeast) at 21-23 °C with a 12h:12h photoperiod.

Heterozygous mutants (*EDTP*^*DJ694*^/+) and their sibling controls (*w*^1118^) were prepared by two consecutive crosses between homozygous *EDTP*^*DJ694*^ mutants and *w*^1118^ flies. We chose virgin female offspring (*EDTP*^*DJ694*^/+) from the first mating and crossed them with *w*^1118^ males. Flies carrying the *EDTP* mutation (red-eyed) and their siblings (white-eyed) were collected for survival analysis.

C57BL/6J mice and Sprague/Dawley rats were purchased from the Experimental Animal Centre of Wuhan University. The C57BL/6J mice are wild-type whereas APP/PS1 (stock number 004462, Jackson Laboratory) are the transgenic mouse models of Alzheimer’s disease. The APP/PS1 mice express a chimeric mouse/human amyloid precursor protein (Mo/HuAPP695swe) and a mutant human presenilin 1 (PS1-dE9) both directed to CNS neurons. The mice were housed in a light/dark (12h:12h) cycle in standard group cages (4-5 per cage) with accessible food and water *ad libitum*. The Sprague/Dawley rats were used for the isolation of primary hippocampal neurons for cell culture. Experimental procedures with mice were approved by the Ethics Committee of Tongji Medical College, Huazhong University of Science and Technology.

### Post-anoxia survival

Newly emerged flies were collected and aged to 6-8 days in regular culture conditions and exposed to an anoxia (generated by pure argon gas) during the light phase of the photoperiod. The percent survival at 12 h (% 12-h survival) after anoxia was examined. Replicated groups (15-25 flies per group, n = 3 ~ 7) of flies were examined. We chose the anoxia exposure of 1, 3, or 6 h because flies are highly tolerant to anoxia for hours [20].

For the tests of post-anoxia long-term survivorship, flies were exposed to anoxia for 0 or 6 h. The dead flies were scored daily during the first week of recovery, and twice a week thereafter until all the flies were counted. Throughout the long-term survival experiments, alive flies were transferred to fresh food vials twice a week.

### Lifespan experiments

Newly emerged flies were collected and raised at a density of 20-25 flies per vial. They were transferred into fresh culture media twice a week. The lifespan was scored by counting dead flies during each transfer until all the flies were counted. This procedure was also used for the examination of the lifespan of flies expressing β amyloid (Aβ) peptides N-42 (Aβ42) or polyglutamine (polyQ) protein aggregates (with 103 tandem repeats) in motoneurons, and the lifespan of flies with simultaneous expression of Aβ42 or polyQ aggregates and RNAi knockdown of *EDTP* in the same cells. Only male flies were tested.

### Cell Culture

The rat primary hippocampal neurons were prepared by following a reported method [21]. Briefly, hippocampal neurons from the 18-day-old embryonic (E18) Sprague/Dawley rats were isolated and seeded at 30,000 - 40,000 cells per well on 6-well plates coated with Poly-D-Lysine/Laminin (Bioscience) in the neurobasal medium (Invitrogen), which was supplemented with 2% B27/0.5 mM glutamine/25 mM glutamate. Half of the culture medium was changed every 3 days with neurobasal medium supplemented with 2% B27 and 0.5 mM glutamine. All cultures were kept at 37 °C in a humidified 5% CO_2_ cultural condition.

Neuro2a cells were cultured at 37 °C in a 1:1 mixture of DMEM (Dulbecco’s modified Eagle’s medium) and OPTI-MEM supplemented with 15% FBS (fetal bovine serum, Gibco), 100 units/ml penicillin, and 100 mg/ml streptomycin. A humidified atmosphere containing 5 % CO_2_ was provided. The cells were plated on to six-well plates overnight and treated with 5 μM Aβ42 or vehicle control (DMSO) for 24 h.

### Immunohistochemistry

Mice were anesthetized and perfused through aorta with 0.9% NaCl followed by phosphate buffer containing 4% paraformaldehyde. The brains were removed and post-fixed in perfusate overnight. Tissue sectioning (20 μm) was performed with a vibratome (Leica Biosystems). The sections were fixed with 0.3% H_2_O_2_ in the absolute methanol for 30 min and saturated with bovine serum albumin (BSA) for 30 min at room temperature. Tissues were then incubated with polyclonal anti-MTMR14 overnight at 4 °C. After wash, the tissues were incubated with appropriate secondary antibody. Immunoreaction was developed with diaminobenzidine using Histostain-SP Kits (Zymed, CA, USA). Images were taken using a light microscope (Olympus BX60, Tokyo, Japan).

### Western Blotting

Western blotting was performed as described previously [22]. The hippocampal region and cortex were quickly removed and homogenized in the sample buffer (Tris-HCl 50mM (pH 7.4), NaCl 150 mM, NaF 10mM, Na_3_VO_4_ 1mM, EDTA 5 mM, benzamidine 2 mM, and phenylmethylsulphonyl fluoride 1 mM). The extracts were mixed with sample buffer (3:1, v/v, containing 200 mM Tris-HCl (pH 7.6), 8% SDS, 40% glycerol, 40mM dithiothreitol) and incubated with boiled water for 10 min. The proteins were separated by 10% SDS-polyacrylamide gel electrophoresis and transferred to nitrocellulose membrane. The membrane was blocked for 1 h with 5% skim milk (dissolved in TBSTween-20 containing 50mM Tris-HCl (pH 7.6), 150mM NaCl, 0.2% Tween-20), and probed with primary antibody at 4 °C overnight. Blots were incubated with anti-rabbit or anti-mouse IgG conjugated to IRDye (800 CW) at room temperature and visualized using the Odyssey Infrared Imaging System (LI-COR Biosciences, Lincoln, NE, USA). Protein bands were analyzed using ImageJ [23].

### Statistics

Statistical analysis was conducted by using R [24] and the following packages: gdata, survival, survminer, and ggplot2. A two-way ANOVA with Tukey’s test was applied for the analysis of 12-h survival of different genotypes of flies which were subject to anoxia with varying durations. A log-rank test was performed for the comparison of two survival curves. A log-rank test with the Benjamini - Hochberg (“BH”) adjustment was used for the comparison of three survival curves. One-way ANOVA with Bonferroni’s multiple comparisons was performed to analyze the MTMR14 expression among three groups of cells. A student *t*-Test was conducted to compare the levels of MTMR14 between two groups of cells. A value of *P* < 0.05 was considered statistical significance.

## Results

### Improved 12-h survival to prolonged anoxia in male flies of *EDTP* mutant

*EDTP*^*DJ694*^ is an enhancer-trap line carrying a transposable element *P{GawB}* in the first intron with the orientation opposite to the affected *EDTP* gene [6,7]. Flies homozygous for *EDTP*^*DJ694*^ are hypomorphic (reduced *EDTP* transcription, short-lived, motor defective, and reduced fecundity) [14]. To avoid the potential confounding effect, we started to examine the flies heterozygous for *EDTP*^*DJ694*^ allele.

*EDTP*^*DJ694*^/+ flies and their sibling controls (*w*^1118^) were raised to 6-8 days old and exposed to an anoxia (generated by pure argon gas) for 1, 3, or 6 h. After 1 h anoxia, the average percentage of survival at 12 h (% 12-h survival) of male flies was 100.0 ± 0.0 % (Mean ± SD) in both *EDTP*^*DJ694*^/+ and controls (Fig. 1a). With 3 h exposure, *EDTP*^*DJ694*^/+ males showed the average % 12-h survival (83.3 ± 10.8 %) higher than controls (67.9 ± 8.4 %) (*P* = 0.0097, Two-way ANOVA with Tukey’s test). After 6 h anoxia, *EDTP*^*DJ694*^/+ males had the % 12-h survival in (63.4 ± 6.7 %) remarkably higher than controls (28.4 ± 5.2 %) (*P* < 0.0001, Two-way ANOVA with Tukey’s test). The genotype, anoxia duration, and their interaction were the factors all significantly contributed to the % 12-h survival in male flies (Two-way ANOVA) (Fig. 1b). Female *EDTP*^*DJ694*^/+ flies had the % 12-h survival statistically the same as *w*^1118^ flies with any of the 1, 3, or 6 h exposure. Two-way ANOVA indicated that while anoxia duration significantly contributed to % 12-h survival, the genotype or the interaction factor had an insignificant contribution (Fig. 1b). Therefore, male *EDTP*^*DJ694*^/+ flies displayed improved 12-h survival to the prolonged anoxia (i.e. 3 - 6 h duration), whereas both *EDTP*^*DJ694*^/+ and control females had comparably enhanced 12-h survival to the prolonged anoxia.

**Figure 1.**
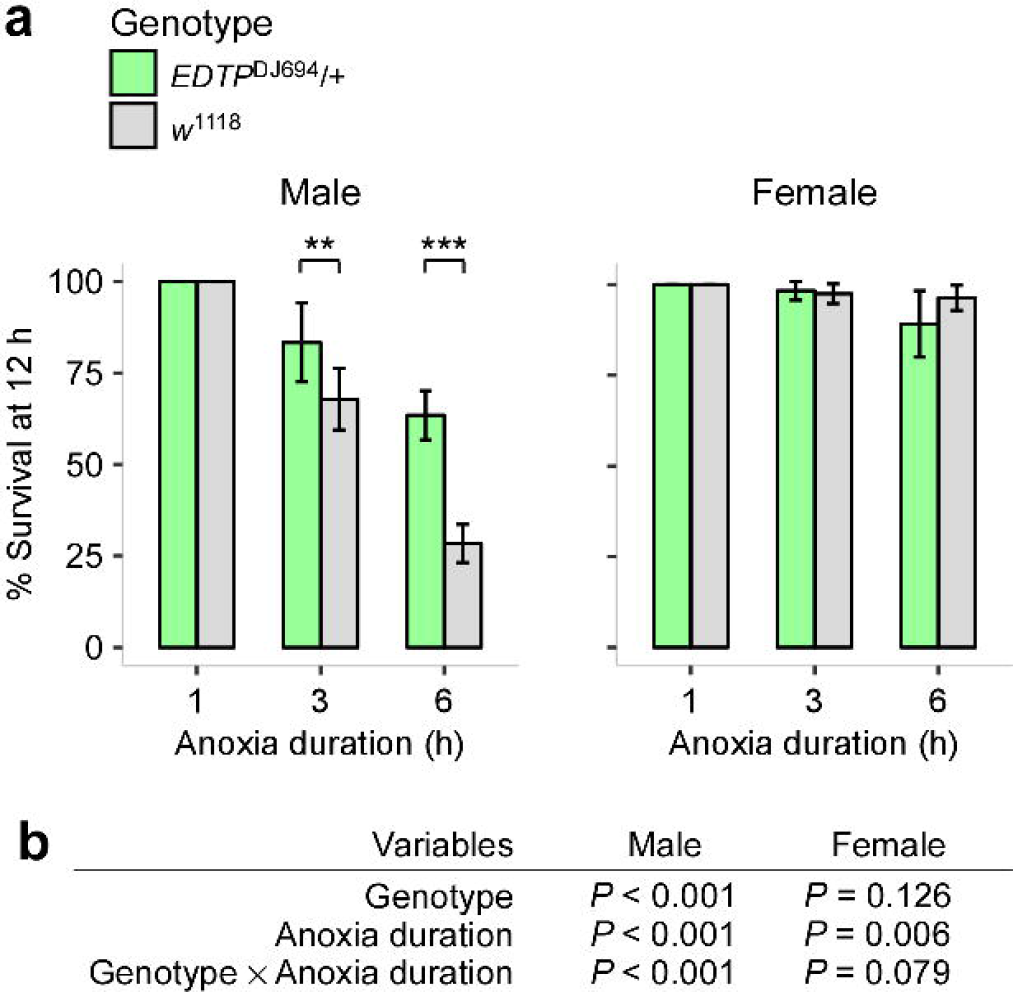
The 12-h survival after anoxia in the *EDTP* mutant. (**a**) % 12-h survival in heterozygous *EDTP* mutants (*EDTP*^*DJ694*^/+, green) and their sibling controls (*w*^1118^, grey) to anoxia. Flies at 6-8 days were exposed to 1, 3, or 6 h anoxia. Left panel, male flies; right panel, female flies; ** *P* < 0.01 and *** *P* < 0.001 from two-way ANOVA with Tukey’s test. (**b**) Two-way ANOVA for the analysis of contributing significance of the genotype, anoxia duration and their interaction to the % 12-h survival.

### Increased post-exposure survivorship to prolonged anoxia in *EDTP* mutant

The increased 12-h survival to prolonged anoxia raised an immediate question: whether *EDTP* mutants have increased post-exposure survivorship throughout the life. Young flies (6-8 days old) were exposed to a 6-h anoxia and the post-exposure survivorship was examined. We used the 6-h exposure because the % 12-h survival was markedly different between heterozygous mutant and controls.

Without anoxia, the median survival time was 69 d (95% confidence interval (CI) 65 - 69 d, n = 102) in *EDTP*^*DJ694*^/+ males and 69 d (CI 65 - 69 d, n = 96) in controls. There was no statistical difference in the survivorship between these groups (*P* = 0.945, Log-rank test) (Fig. 2). There was also no significant difference in the survivorship between female *EDTP*^*DJ694*^/+ (median 74 d, CI 69 - 79 d, n = 107) and their controls (median 74 d, CI 69 - 79 d, n = 97) (*P* = 0.869, Log-rank test). Thus, without anoxia, the *EDTP* mutation displayed no effect on the survivorship.

**Figure 2.**
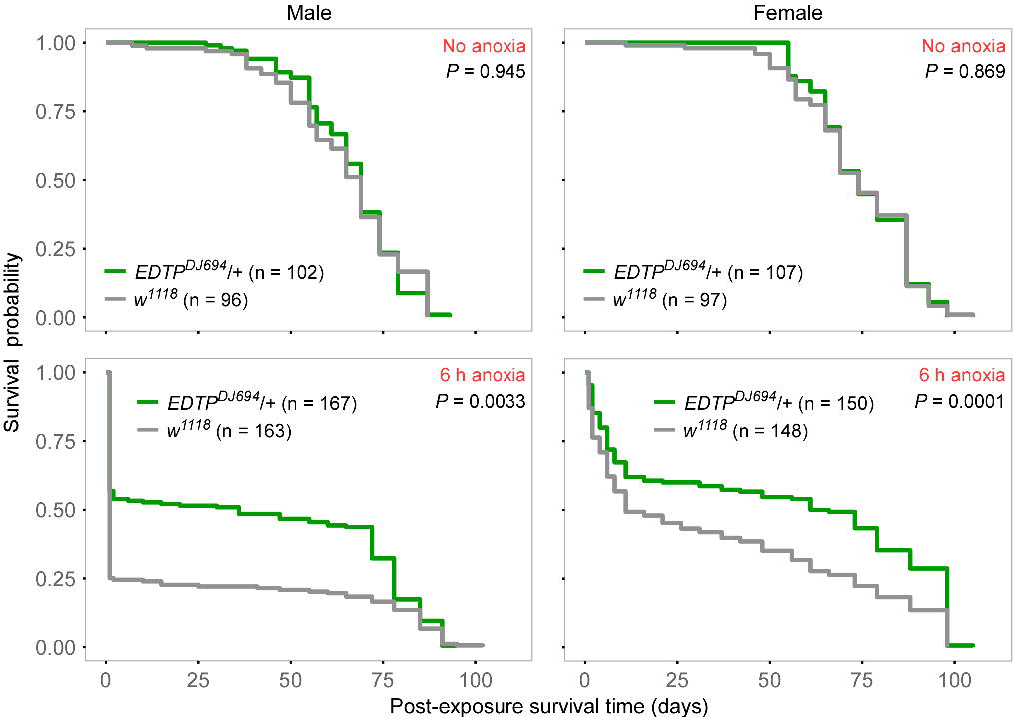
Improved survivorship after prolonged anoxia in the heterozygous *EDTP* mutant. Flies (6-8 days old) were exposed to 0 or 6 h anoxia. The post-exposure survival curves of flies carrying *EDTP* mutation (*EDTP*^*DJ694*^/+, green) and their sibling controls (*w*^1118^, grey) were plotted. Anoxia durations (red text) are indicated. *P* values are from Log-rank test. Left panels, male flies; right panels, female flies.

With 6 h anoxia, *EDTP*^*DJ694*^/+ males showed post-exposure survival (median 36 d, CI 1 - 72 d, n = 167) markedly longer than controls (median 1 d, CI 1 - 1 d, n = 163) (*P* = 0.0033, Log-rank test). *EDTP*^*DJ694*^/+ females also showed the post-exposure survival (median 63.5 d, CI 37 - 79 d, n = 150) remarkably longer than controls (median 11 d, CI 8 - 37 d, n = 148) (*P* = 0.0001, Log-rank test). Both male and female flies of *EDTP*^*DJ694*^/+ had increased post-exposure survivorship to 6 h anoxia.

### RNAi knockdown of *EDTP* in motoneurons extended lifespan

Extreme hypoxia induces autophagy in mammalian and human cells [18]. The mammalian homologous *MTMR14* negatively regulates the autophagic removal of protein aggregates [11]. These data together with our findings suggested that downregulation of *EDTP* could promote autophagy and improve the survivorship. We used the Gal4/UAS binary system [17] for selective downregulation of *EDTP* in motoneurons and examined the lifespan. The rationale for targeting motoneurons was to avoid affecting the abundant expression of EDTP in muscles, ovaries, spermatheca, and oocytes. The motoneuron-specific driver D42-Gal4 [25] was used for RNAi knockdown.

We used the UAS-EDTP-RNAi (III) line with a short interference sequence targeting the transcripts from exon 2 of *EDTP*. Flies with *EDTP* knockdown in motoneurons (D42-Gal4/UAS-EDTP-RNAi (III)) had a median lifespan of 113 d (CI 107 – 118 d, n = 132), which was longer than UAS controls (UAS-EDTP-RNAi (III)/+, median 89 d, CI 89 – 98 d, n = 63) (*P* < 0.0001, Log-rank test with “BH” adjustment), and also longer than Gal4 controls (D42-Gal4/+, median 73 d, CI 73 – 77 d, n = 137) (*P* < 0.0001, Log-rank test with “BH” adjustment) (Fig. 3a). Therefore, downregulation of *EDTP* in motoneurons resulted in extended lifespan.

**Figure 3.**
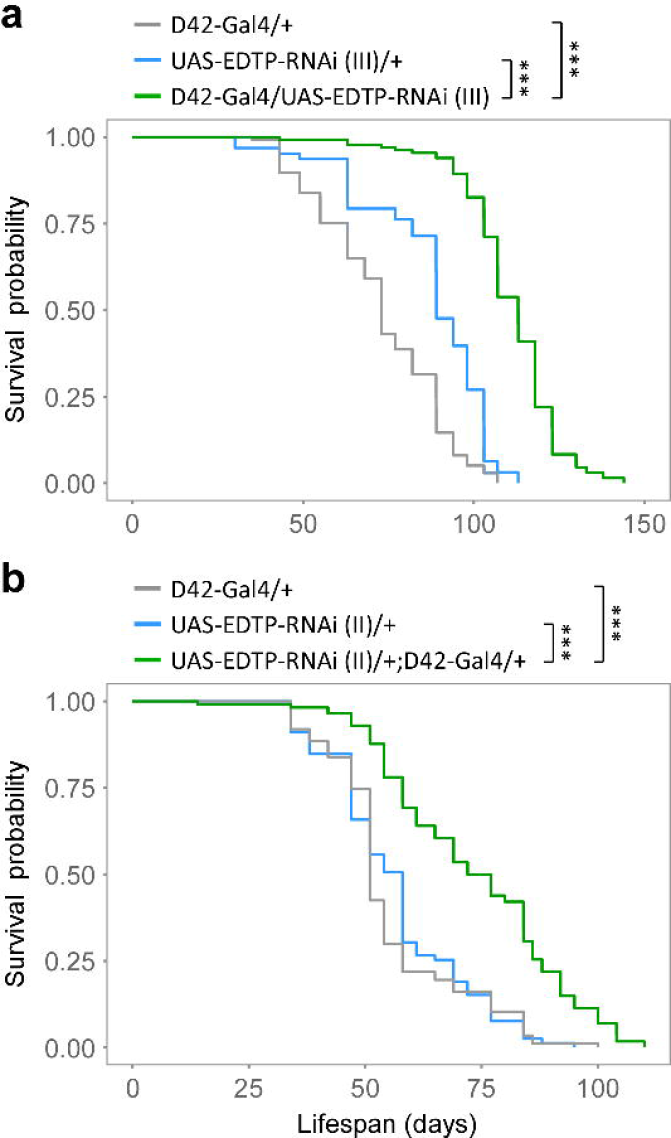
Downregulation of *EDTP* in motoneurons extended lifespan. (**a**) The survival curves of flies with *EDTP* knockdown (D42-Gal4/UAS-EDTP-RNAi (III), green) and their controls (D42-Gal4/+, grey; UAS-EDTP-RNAi (III)/+, light blue). (**b**) The survival curves of flies with *EDTP* knockdown (UAS-EDTP-RNAi (II)/+; D42-Gal4/+, green) and their controls (D42-Gal4/+, grey; UAS-EDTP-RNAi (II)/+, blue). Two UAS lines carrying independent RNAi constructs, one in the third chromosome and another in the second chromosome, were used. Presented data are from male flies. *** *P* < 0.001 from Log-rank test with “BH” adjustment.

To confirm this finding, we used a different RNAi line (UAS-EDTP-RNAi (II) with an interference sequence targeting the transcripts from exon 3). Flies with *EDTP* knockdown in motoneurons (UAS-EDTP-RNAi (II)/+; D42-Gal4/+) had the lifespan (median 74.5 d, CI 69 – 84 d, n = 114) significantly longer than UAS controls (median 58 d, CI 51 – 58 d, n = 79) (*P* < 0.0001, Log-rank test with “BH” adjustment), or Gal4 controls (median 51 d, CI 51 – 54 d, n = 87) (*P* < 0.0001, Log-rank test with “BH” adjustment) (Fig. 3b). These data indicated that downregulation of *EDTP* in motoneurons extended the lifespan in *Drosophila*.

### *EDTP* knockdown in motoneurons improved the survivorship to beta-amyloid peptides or polyglutamine protein aggregates

We next asked whether downregulation of *EDTP* could increase the survivorship to cellular protein aggregates expressed in the motoneurons. We examined the survivorship of flies with simultaneous *EDTP* downregulation and the expression of toxic β amyloid (Aβ) peptides N-42 (Aβ42) [26]. The median lifespan in flies expressing Aβ42 in motoneurons (UAS-Aβ42/Y;; D42-Gal4/+) was 64 d (CI 64 - 64 d, n = 111). The median lifespan was increased to 94 d (CI 94 - 98 d, n = 86) in flies with Aβ42 expression and simultaneous RNAi knockdown of *EDTP* (UAS-Aβ42/Y;; D42-Gal4/UAS-EDTP-RNAi) (*P* < 0.0001, Log-rank test) (Fig. 4a).

**Figure 4.**
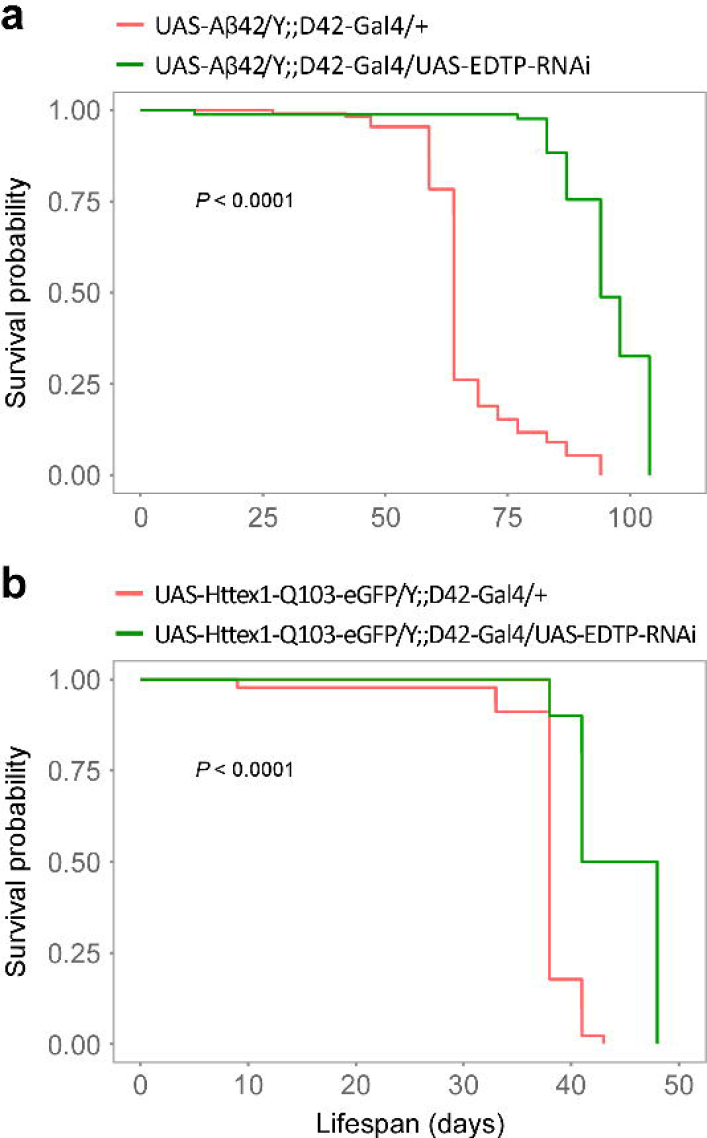
Downregulation of *EDTP* in motoneurons improved the survivorship to the protein aggregates expressed in the same cells. (**a**) The survival curves of flies expressing Aβ42 (red) and flies with both Aβ42 expression and *EDTP* knockdown in the motoneurons (green). *P* value from a log-rank test. (**b**) The survival curves of flies expressing polyQ protein aggregates (red) and flies with both polyQ expression and *EDTP* knockdown in the motoneurons (green). *P* value from a log-rank test. Notes: D42-Gal4, motoneuron-specific driver; UAS-Aβ42, a UAS line carrying a gene encoding human beta-amyloid peptide (N = 42); UAS-Httex1-Q103-eGFP, a UAS line for the expression of polyQ protein aggregates (containing in each molecule a polyglutamine tract with 103 tandem repeats); UAS-EDTP-RNAi, an RNAi line (#36917) carrying an interference sequence targeting the exon 2 of *EDTP*. Data are from male flies.

We also examined the survivorship of flies with *EDTP* downregulation and simultaneous expression of polyglutamine (polyQ) protein aggregates in the motoneurons. The median lifespan was 38 d (CI 38 - 38 d, n = 45) in flies (UAS-Httex1-Q103-eGFP/Y;; D42-Gal4/+) expressing polyQ aggregates (each containing a polyQ tract of 103 tandem repeats of glutamine). The median lifespan was significantly increased to 44.5 d (CI 41 – 48 d, n = 30) in flies (UAS-Httex1-Q103-eGFP/Y;; D42-Gal4/UAS-EDTP-RNAi) with both polyQ expression and *EDTP* downregulation in motoneurons (*P* < 0.0001, Log-rank test) (Fig. 4b).

Therefore, *EDTP* downregulation in motoneurons improved the survivorship to the expression of Aβ42 or polyQ protein aggregates.

### Expression of MTMR14 in the hippocampus and cortex in C57BL/6J and APP/PS1 mice

Human *MTMR14* is transcribed ubiquitously at relatively low levels in the brain [27]. Mouse MTMR14 is expressed in the muscles, liver, and fat but there is little evidence for the brain expression [28]. We examined the expression of MTMR14 in the hippocampus and cortex in C57BL/6J and APP/PS1 mice. C57BL/6J is wild-type, whereas APP/PS1 is a transgenic strain with constitutive expression of a chimeric mouse/human amyloid precursor protein (Mo/HuAPP695swe) and a mutant human presenilin 1 (PS1-dE9) in CNS neurons [29].

MTMR14 expression was observable in the CA3 pyramidal neurons and dentate gyrus (DG) in the hippocampus of a 3-month-old C57BL/6J mouse (Fig. 5a). MTMR14 expression was detectable in the hippocampus in both C57BL/6J and APP/PS1 mice. The levels in two strains were similar with no significant difference (Fig. 5b). MTMR14 expression was also detectable in the cortex in both strains. However, relative levels of MTMR14 in APP/PS1 mice were lower than those in C57BL/6J mice (*P* < 0.05, *t*-Test). These data indicated an association between the transgenic expression of amyloid precursor protein/presenilin 1 and MTMR14 level.

**Figure 5.**
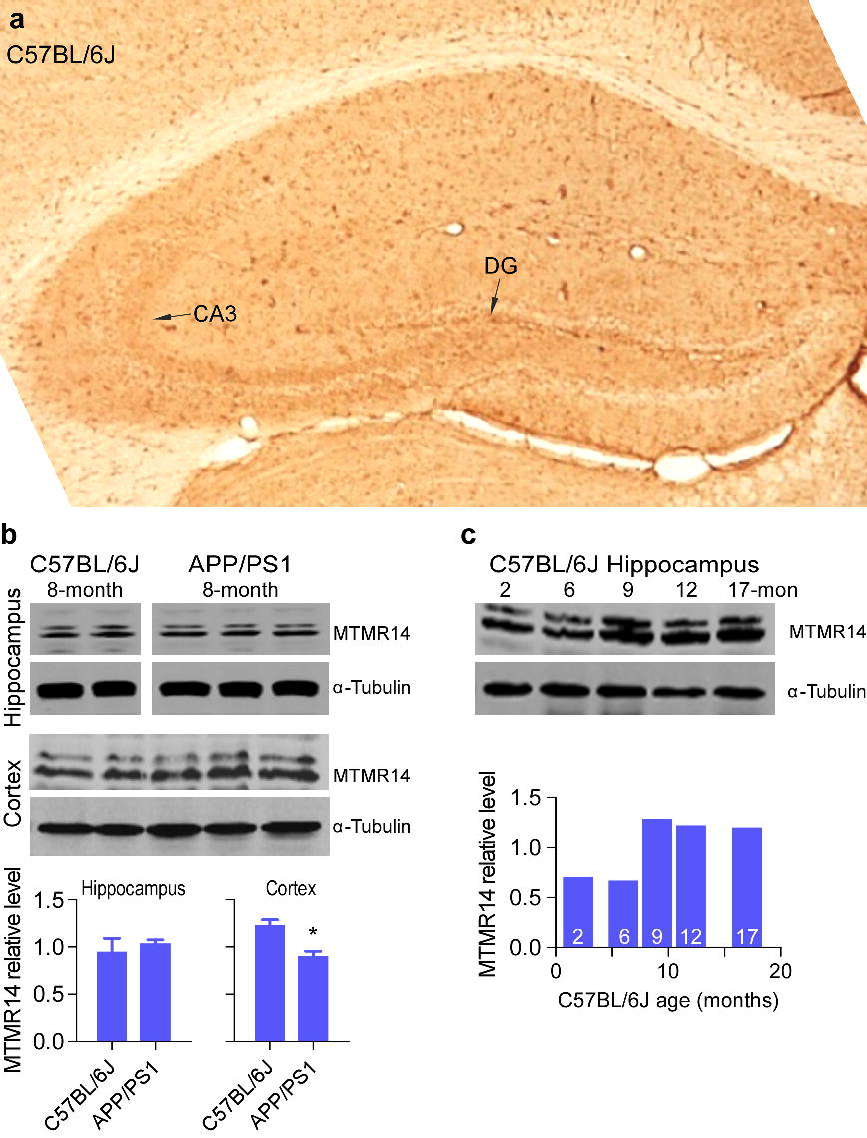
Expression of MTMR14 in the hippocampus and cortex in C57BL/6J and APP/PS1 mice. (**a**) Immunostaining of MTMR14 in the hippocampus in a C57BL/6J mouse. CA3, CA3 pyramidal neurons; DG, dentate gyrus. (**b**) Expression of MTMR14 in the hippocampus and cortex in C57BL/6J and APP/PS1 mice. α-Tubulin, loading control; * *P* < 0.05, Student’s *t*-test. (c) Expression of MTMR14 in the hippocampus in C57BL/6J mice at different ages (in months). Numbers in the bars indicate the ages of mice.

Relative levels of MTMR14 in the C57BL/6J hippocampus increased and plateaued at 9, 12 and 17 months compared with the levels at 2 and 6 months (Fig. 5c), suggesting an age-dependent kinetics of MTMR14 expression.

### Increased MTMR14 induced by Aβ42 in the rat primary hippocampal neurons and Neuro2a cells

Aβ42 causes microRNA deregulation in primary hippocampal neurons in C57BL/6J mice [30] and inhibits the viability of the mouse neuroblastoma Neuro2a cells [31]. We explored the effect of Aβ42 treatment on MTMR14 expression in primary hippocampal neurons and Neuro2a cells.

The primary hippocampal neurons from 18-day-old embryonic (E18) Sprague/Dawley rats were prepared and treated with Aβ42. Relative MTMR14 levels were increased in the primary cultures after a 72-h incubation with 10 μM Aβ42 compared with controls (*P* < 0.01, One-way ANOVA with Bonferroni’s multiple comparisons) (Fig. 6a). There was no significant increase of relative MTMR14 levels in the cultures incubated with 5 μM Aβ42 for 72 h. Data indicated a dosage-dependent induction of MTMR14 by Aβ42 treatment. We examined the MTMR14 expression in Neuro2a cells with Aβ42 treatment. MTMR14 expression was increased in Neuro2a cells treated with Aβ42 at 5 μM for 24 h compared with controls (*P* < 0.01, *t*-Test) (Fig. 6b). Therefore, Aβ42 treatment increased the expression of MTMR14 in both the primary hippocampal neurons of rats and Neuro2a cells.

**Figure 6.**
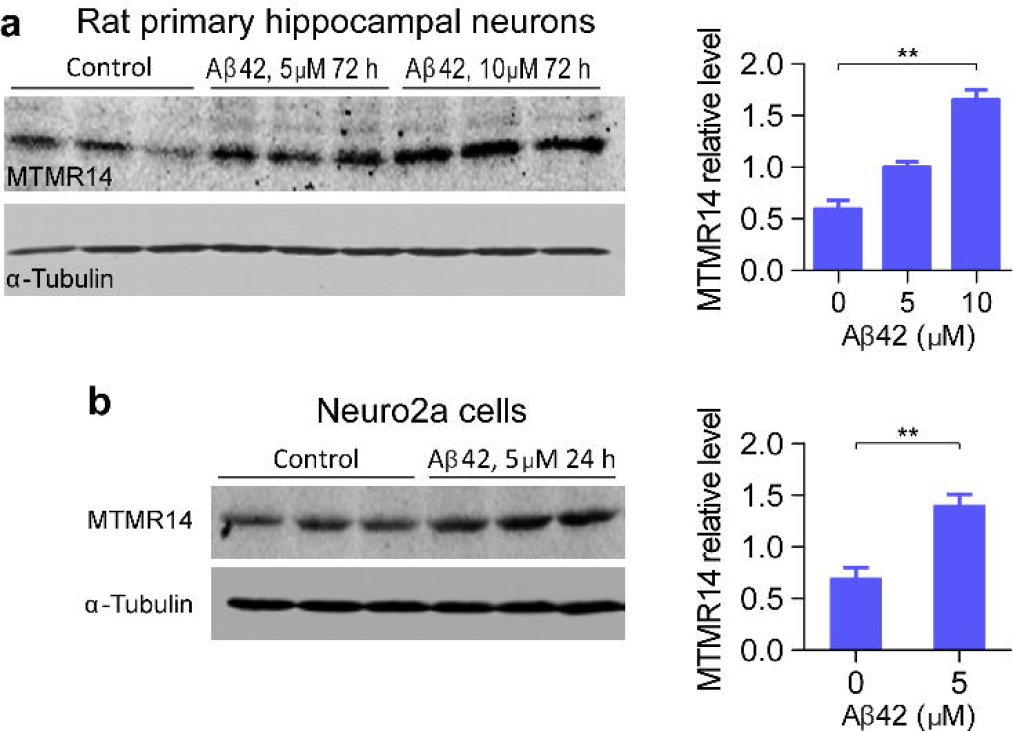
MTMR14 expression induced by Aβ42 in the rat primary hippocampal neurons and Neuro2a cells. (**a**) MTMR14 induction by Aβ42 treatment (0, 5, and 10 μM) for 72 h in the rat primary hippocampal neurons. ** *P* < 0.01 by one-way ANOVA and Bonferroni’s multiple comparisons. (**b**) MTMR14 expression induced by Aβ42 treatment (0 and 5 μM) for 24 h in Neuro2a cells. ** *P* < 0.01, Student’s *t*-test.

## Discussion

We report in *Drosophila* a novel approach of fine-tuning the disease-associated gene *EDTP/MTMR14* for lifespan extension and improved survivorship to cellular protein aggregates. Cell-specific downregulation of *EDTP* in non-muscle tissues circumvents the deleterious consequences of ubiquitous loss of *EDTP*. Specifically, downregulation of *EDTP* in motoneurons extends lifespan and increases the survivorship to Aβ42 or polyQ protein aggregates. The expression of the mouse *MTMR14* in the hippocampus and cortex, together with its age-dependent kinetics and the Aβ42-induced increase of MTMR14 in the rat’s cultured hippocampal neurons, promises a potential application by targeting *EDTP/MTMR14* in a tissue-specific manner for the treatment of neurodegeneration.

The proposed cell-specific downregulation of *EDTP/MTMR14* for beneficial consequences is formulated largely from the preliminary observation that heterozygous flies carrying the *EDTP*^*DJ694*^ allele have increased survivorship to prolonged anoxia. Homozygous flies carrying this allele display a “jumpy” phenotype – impaired motor function associated with shortened lifespan and reduced fecundity [14]. These findings indicate that downregulation of *EDTP* in flies homozygous for *EDTP*^*DJ694*^ is overall deleterious, whereas moderate downregulation of *EDTP* in heterozygous flies causes no obvious motor defect and becomes protective if flies are exposed to a prolonged anoxia. Such a conditional benefit highlights the importance of subtle manipulation of spatiotemporal expression of EDTP/MTMR14 for the desired effect. *EDTP/MTMR14* has two main functions: negative regulation of autophagy and disease-causing effect if deficient in the muscles [1,11–15]. *Drosophila EDTP* has two peaks of expression, one at oogenesis and another at adult stages around 20-30 days [4–7]. Both male and female adult flies have abundant EDTP in muscles and additionally, female flies express rich amounts of EDTP in the spermatheca and ovaries. The differential expression of EDTP in female flies might also be responsible for the enhanced short-term survival (i.e. % 12-h survival) to prolonged anoxia. The dual functions, multiple peaks over time, broad expressing tissues, and the sexual dimorphism of expression pattern together do not likely allow a complete deletion of *EDTP/MTMR14* to be associated with beneficial consequences. Precise control of EDTP/MTMR14 expression in the favorable cells or tissues could be essential for eliciting the protective effects. The current study presents a strategy of cell-specific downregulation of *Drosophila EDTP* in motoneurons with beneficial effects in lifespan extension and improved survivorship to cellular protein aggregates.

Motoneurons are highly specialized and terminally differentiated with reduced function of global genome repair [32,33]. More importantly, neuronal cells utilize autophagy as an essential process for normal turnover of cytoplasmic contents [34]. *Drosophila* motoneuronal expression of a human Cu-Zn superoxide dismutase (SOD1) leads to a marked extension of lifespan by up to 40% [35]. Motoneuronal overexpression of the heat shock protein 70 results in structural plasticity of axonal terminals which is associated with increased larval thermotolerance [36]. Motoneurons innervate muscle cells through neuromuscular junctions, making them an ideal target for *EDTP* downregulation while leaving muscular expression intact. The findings that motoneuronal downregulation of *EDTP* extends lifespan and improves the survivorship to Aβ42 or polyQ protein aggregates firmly support motoneurons as a favorable targeting tissue. Notably, our findings greatly rely on the putative motoneuronal driver D42-Gal4, which also shows a prominent expression pattern in the peripheral sensory neurons [37]. Whether sensory neurons utilize the EDTP/MTMR14-associated autophagy for removing cellular wastes is unclear, but a study has shown the connection between the sensory neurons and lifespan extension [38].

EDTP/MTMR14 possesses a function opposite to Vps34 in the regulation of the PtdIns3P pool. Vps34 is the only phosphatidylinositol 3-kinases in yeast [2] and has been evolutionarily conserved through mammals. Vps34 plays an essential role in the process of autophagy [39,40]. Downregulation of *EDTP/MTMR14* shifts the focus from Vps34 to its functional opponent for the regulation of PtdIns3P and thus represents a novel approach to manipulate the process of autophagy.

The evident expression of MTMR14 in the mouse hippocampus and cortex provides a foundation from which the cell-specific manipulation of *EDTP/MTMR14* expression could be applied in mammals. Furthermore, the transgenic expression of a chimeric mouse/human amyloid precursor protein (Mo/HuAPP695swe) and a mutant human presenilin 1 (PS1-dE9) in APP/PS1 mice is associated with the downregulation of MTMR14 in the cortex but not in the hippocampus. These findings indicate an involvement of MTMR14 expression in the deposit of amyloid peptides and this involvement is tissue-specific. Moreover, MTMR14 expression displays an age-related kinetics and MTMR14 is inducible by Aβ42 treatment in the rat primary hippocampal neurons as well as mouse Neuro2a cells, suggesting a close relation between MTMR14 expression and aging. Noteworthy, MTMR14 expression in response to the presence of amyloid peptides is different between *in vivo* animal experiments and cultured neurons. Perhaps the increase of MTMR14 in cultured neurons represents a short-term response to Aβ42, whereas MTMR14 downregulation in the animal studies is involved in an active process for the removal of amyloid peptides. Nevertheless, our findings highlight a promising application through targeting EDTP/MTMR14 for lifespan extension and survival improvement to cellular protein aggregates in mammals.

## Acknowledgments

This study was funded by the Fundamental Research Funds for the Central Universities from Nanjing University of Science and Technology (30918011308) and Natural Science Foundation of China (81501211 to D.J. L.).

